# A dual sgRNA approach for functional genomics in *Arabidopsis thaliana* ^[OPEN]^

**DOI:** 10.1101/172676

**Authors:** Laurens Pauwels, Rebecca De Clercq, Jonas Goossens, Sabrina Iñigo, Clara Williams, Mily Ron, Anne Britt, Alain Goossens

**Author notes:** **Corresponding author:** Laurens Pauwels, Department of Plant Systems Biology, VIB-Ghent University, Technologiepark 927, B-9052 Gent (Belgium).

## Abstract

Reverse genetics uses loss-of-function alleles to interrogate gene function. The advent of CRISPR/Cas9-based gene editing now allows to generate knock-out alleles for any gene and entire gene families. Even in the model plant *Arabidopsis thaliana*, gene editing is welcomed as T-DNA insertion lines do not always generate null alleles. Here, we show efficient generation of heritable mutations in Arabidopsis using CRISPR/Cas9 with a workload similar to generating overexpression lines. We obtain Cas9 null-segregants with bi-allelic mutations in the T2 generation. Out of seven new mutant alleles we report here, one allele for *GRXS17*, the ortholog of human GRX3/PICOT, did not phenocopy previously characterized nulls. Notwithstanding, the mutation caused a frameshift and triggered nonsense-mediated decay. We demonstrate that our workflow is also compatible with a dual sgRNA approach in which a gene is targeted by two sgRNAs simultaneously. This paired nuclease method can result in a more reliable loss-of-function alleles that lack a large essential part of the gene. The ease in the CRISPR/Cas9 workflow should help in the eventual generation of true null alleles of every gene in the Arabidopsis genome, which will advance both basic and applied plant research.

**One-sentence summary:** We present a dual sgRNA approach to delete Arabidopsis gene 34 fragments in order to obtain reliable functional knock-outs.

## INTRODUCTION

The precise introduction of a DNA double-strand break (DSB) in a plant genome can now be accomplished by a variety of techniques (Baltes and Voytas, 2015). However, the advent of Clustered Regularly Interspaced Short Palindromic Repeats/CRISPR associated 9 (CRISPR/Cas9)-based technology has brought reliable gene editing (GE) within the reach of non-specialized molecular biology labs. The power of CRISPR/Cas9 compared to predecessor techniques lies in both its consistent high efficiency and its simple two-component design. A generic nuclease, Cas9, is guided to a target DNA sequence (protospacer) by associating with an artificial single guide RNA (sgRNA) (Jinek *et al.*, 2012). Changing the typically 20 nucleotide long target-specific spacer sequence in the sgRNA is sufficient for redirecting the RNA-guided engineered nuclease to another genomic locus. In addition, several sgRNAs with different targets can be co-expressed allowing for multiplexing as exemplified in *Arabidopsis thaliana* by targeting of the PYRABACTIN RESISTANCE1-LIKE (PYL) family of abscisic acid receptor genes (Zhang *et al.*, 2016) or the GOLVEN family (Petserson *et al.*, 2016).

DSBs are readily recognized by the plant cell and repaired. When non-homologous end-joining (NHEJ) pathway results in imprecise repair, small insertions and deletions (indels) at the cut sit are produced (Knoll *et al.*, 2014). In Arabidopsis, 1 base pair (bp) insertions (+1) are usually observed (Fauser *et al.* 2014, Feng *et al.*, 2014). The alternative NHEJ (aNHEJ) pathway uses a molecularly distinct mechanism and microhomologies flanking the cut site guide repair. Also known as microhomology-mediated end joining (MMEJ), it results in relatively larger deletions (Knoll *et al.*, 2014). NHEJ-mediated indel-formation is used to generate loss-of-function mutants. If the indel causes a frame-shift, a non-functional truncated protein can be translated and/or a premature stop codon will trigger nonsense-mediated decay (NMD) causing organized mRNA degradation by the cell (Popp and Maquat, 2016) CRISPR/Cas9 technology has been established for Arabidopsis and is continuously being developed further (Feng *et al.* 2013, Mao *et al.* 2013, Fauser *et al.* 2014, Feng *et al.*, 2014, Ma *et al.*, 2015, Wang *et al.* 2015, Osakabe *et al.* 2016, Tsutsui and Higashiyama, 2016, Zhang *et al*, 2016, Denbow *et al.*, 2017, Peterson *et al.*, 2016). Reports using CRISPR/Cas9 in Arabidopsis are emerging that are not technology-focused, but rather limited in number taking into account the widespread use of this model organism, the short generation time and its ease of transformation (Gao *et al.*, 2015, Ning *et al.*, 2015, Xin *et al.*, 2016, Zhang *et al.*, 2016, Guseman *et al.*, 2017, Li *et al.*, 2017, Lu *et al.*, 2017, Ritter *et al.*, 2017). The difficulties of using CRISPR/Cas9 to generate mutants in Arabidopsis have been attributed to the unique floral dip system of transformation in which inflorescences of T0 plants are infected with *Agrobacterium tumefaciens*. Primary transformants (T1) are derived via this process from a transformed egg cell (Bechtold *et al.*, 2000). Chimerism, i.e. the presence of at least 3 different alleles, points to Cas9 activity at later stages during somatic growth. This indicates that the mutation did not occur within the egg cell or zygote, but rather after the first cell division. Furthermore, even when mutations are detected in T1 somatic cells, often WT alleles are retrieved when the CRISPR/Cas9 T-DNA has been segregated away (Wang *et al.*, 2015). This can be attributed to gene editing efficiency, i.e. the percentage of cells not WT, as the limited amount of cells that make up the germ line have to be mutated for heritability.

Here, we report and quantify high editing efficiencies in T1 somatic cells and inheritance of NHEJ-repaired alleles in Arabidopsis. Our workflow allows us to obtain Cas9 null-segregants with bi-allelic mutations in the T2 generation. Moreover, it is compatible with a dual sgRNA approach, leading to deletion of gene fragments and greater confidence in loss-of-function alleles.

## RESULTS

### High gene editing efficiency in T1 somatic tissue

The vector pDE-Cas9 has successfully been used for gene editing (GE) in Arabidopsis (Fauser *et al.*, 2014). It contains an Arabidopsis codon-optimized SpCas9 sequence, driven by the *Petroselinum crispum* Ubiquitin4-2 promoter (pPcUBI). As kanamycin resistance is more often used in our lab both in Arabidopsis and in tomato, we used pDE-Cas9Km (Ritter *et al.*, 2017) in which the basta resistance cassette in pDE-Cas9 is replaced with *nptII* (Figure S1). In order to evaluate these vectors, we initially designed nine sgRNAs targeting five genes of interest: *JASMONATE ASSOCIATED MYC2 LIKE 2* (*JAM2*, Sasaki-Sekimoto *et al.*, 2013), *VQ19* and *VQ33* (Jing and Lin, 2015), *HEMOGLOBIN 3* (*GLB3*) and *GLUTAREDOXIN S17* (*GRXS17*) (Nagels Durand *et al.*, 2016). sgRNAs were designed to minimize possible off-target activity (Lei *et al.*, 2014), and when possible predicted sgRNA efficiencies were taken into account (Chari *et al.*, 2015). An updated overview of estimated sgRNA parameters by CRISP-OR (http://crispor.tefor.net/, Haeussler *et al.*, 2016) can be found in Table 1. Although it is currently unknown if the models for sgRNA efficiency, based on empirical data from metazoan cells holds true in plants, we anticipate that at least some sgRNA sequence parameters will be the similar as CRISPR/Cas9 is a fully heterologous system. Preferably sgRNAs were chosen in the 5’ end of the first exon (Figure 1). In the case of *JAM2*, we specifically designed two sgRNAs that targeted the sequence encoding the JAZ interaction domain (JID) (Fernandez-Calvo *et al.*, 2009).

**Figure 1.**
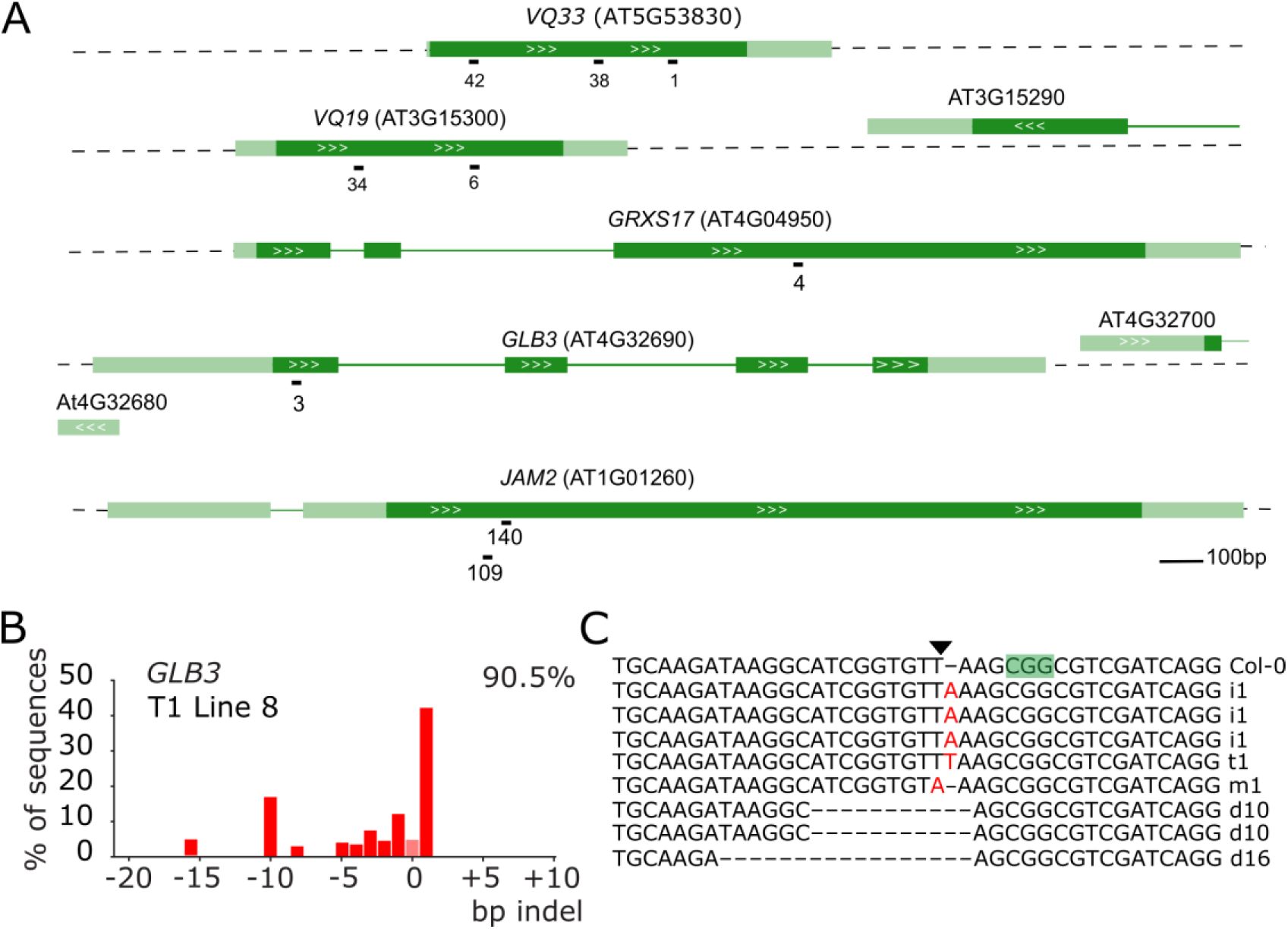
Use of TIDE to analyze CRISPR/Cas9-induced somatic mutations in T1 Arabidopsis plants. A, genomic structure of the targeted genes and location of the sgRNAs. Dark green boxes designate exons; light green boxes, UTRs; solid lines, introns; white arrows gene orientation. sgRNA numbers are arbitrary identifiers. B, example result of a TIDE analysis. A leaf of a T1 plant expressing a CRISPR/Cas9 construct targeting *GLB3* was used to prepare genomic DNA. The targeted region was amplified by PCR and sequenced using standard Sanger sequencing. TIDE software was used to visualize the indel spectrum and estimate overall editing efficiency (top right corner). Bars indicate the number of sequences with a given indel size. Pink bar (indel size of zero) represents WT or base substitution alleles. C, Verification of TIDE using sequencing of individual amplicon subclones. The PAM is highlighted in green, the triangle points to the Cas9 cut site. I, insertion, d, deletion, m, mutation are followed with the number of bases involved.

**Figure S1.**
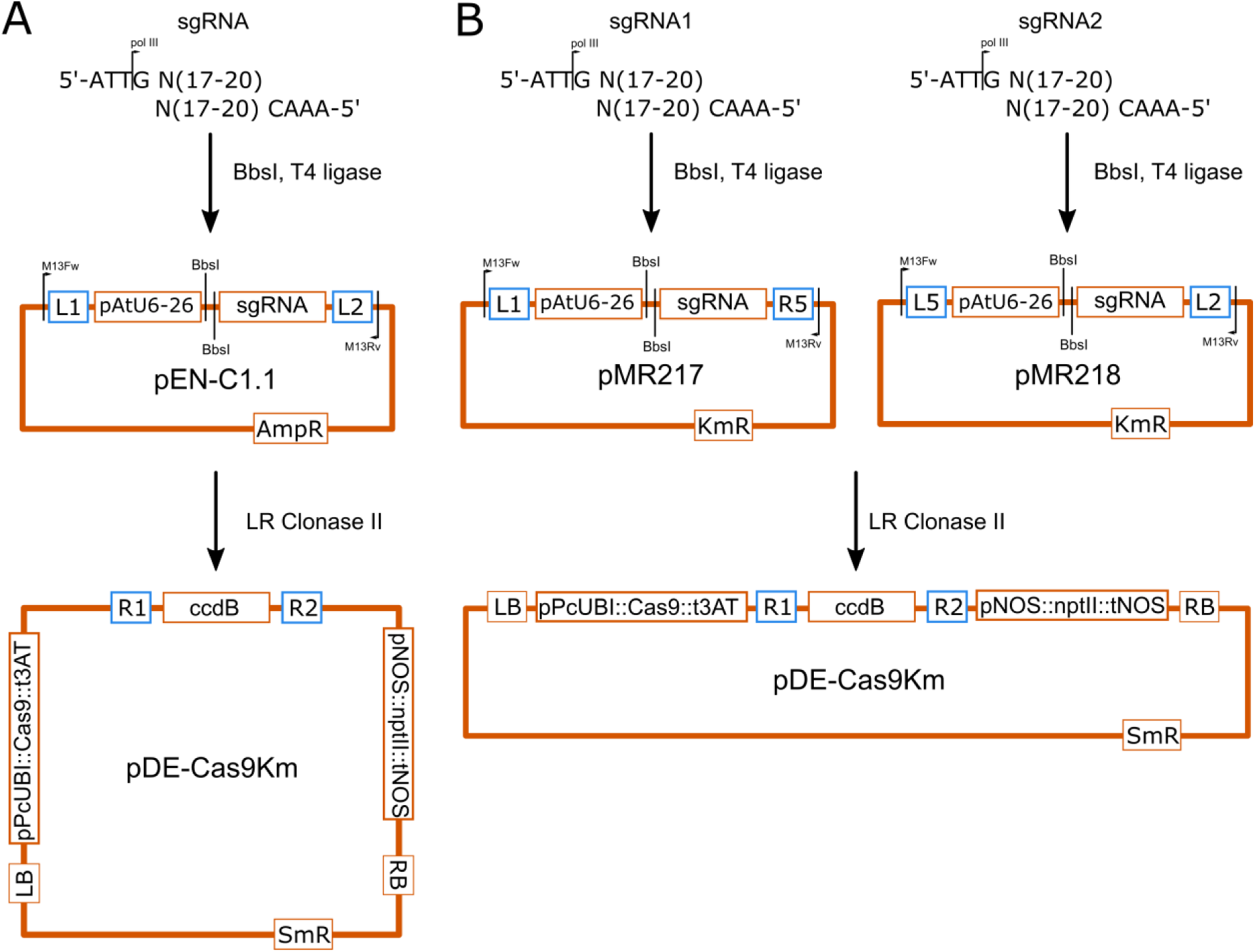
Cloning procedures and vector maps. A, procedure for cloning one sgRNA according to Fauser *et al.*, 2014. Two oligonucleotides are synthesized that have 4-bp overhangs for Type-IIS cloning and contain a guide sequence of 17 to 20 nucleotides. The annealed oligos are cloned in the pEN-C1.1 shuttle vector that contain the Arabidopsis U6-26 promoter driving the sgRNA. The G of the 5’ ATTG overhang is the first transcribed base by the RNA polymerase III. After sequence verification, the vectors are recombined with the pDE-Cas9Km destination vector that contains the Cas9 (codon optimized for *Arabidopsis thalaiana*) under control of the *Petroselinum crispum* ubiquitin4-2 promoter (pPcUBI) and the *nptII* selection marker for plants B, dual sgRNA cloning. The pMR217 or pMR218 shuttle vectors used are identical, except for the MultiSite Gateway recombination sites (in blue). AmpR, KmR and SmR are ampicillin, kanamycin and streptomycin resistance markers for *Escherichia coli*.

The sgRNA cloning procedure (Figure S1A) uses the type II restriction enzyme BbsI and utilizes a 5’ ATTG overhang of which the G serves as the first nucleotide of the sgRNA when transcribed by the polymerase III promoter AtU6-26. Most sgRNAs were of the GN19-type with the 5’ G being the first transcribed base of a 20-bp long guide sequence. One sgRNA, JAM2-140, was of the GN20-type. An extra 5’ G or GG attached to the sgRNA should not hinder efficiency (Cho *et al.*, 2014). Another sgRNA VQ33-42, was a GN18-type. Truncated sgRNAs (tru-gRNAs) down to a 17bp guide sequence have been shown to be as efficient as 20bp guides in human cells (Fu *et al.*, 2014).

For each single sgRNA construct, approximately 15 T1 Arabidopsis plants were selected on basta or kanamycin respectively. One of the first true leaves was harvested for genomic DNA extraction. A region spanning the predicted cut site was amplified by PCR and the amplicon sequenced by traditional Sanger sequencing. Arabidopsis CRISPR/Cas9 T1 plants are typically chimeric, defined as having at least three different alleles for a locus (Feng *et al.*, 2014). Different cell files showed different indels in both alleles after NHEJ-mediated repair, leading to a range of detectable indels in a single leaf and a complex chromatogram. The quantitative sequence trace data was therefore decomposed using the Tracking of Indels by DEcomposition (TIDE) software (https://tide.nki.nl/) (Brinkman *et al.*, 2014). This results in an estimation of overall editing efficiency (percentage of cells not WT) and the spectrum and frequency of the dominant indel types (See Figure 1B for an example for *GLB3*). Subcloning of amplicons followed by sequencing yielded similar profiles (Figure 1C). Furthermore, examination of genomic DNA of different leaves yielded comparable but not identical patterns (Figure S2).

**Figure S2.**
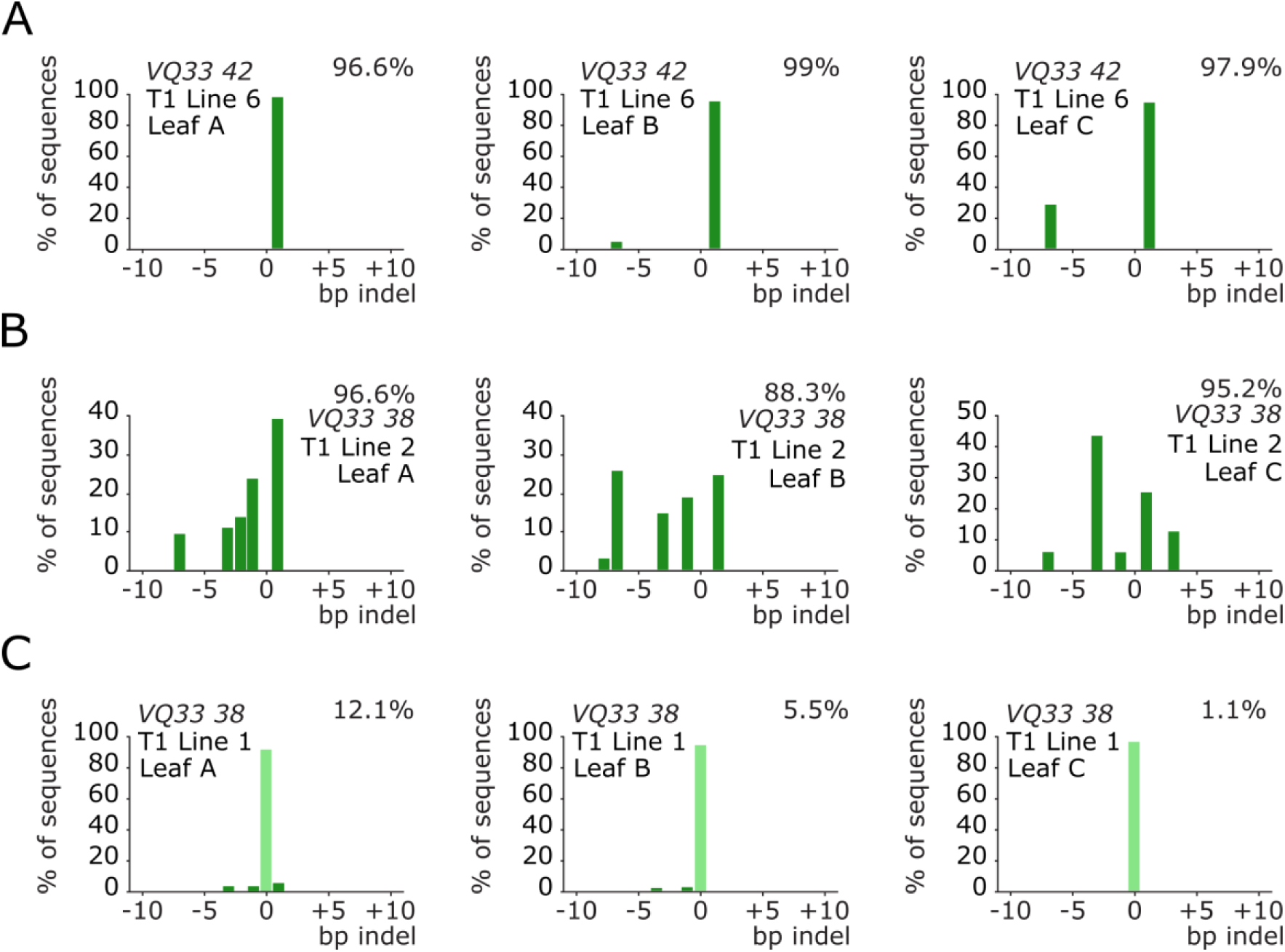
Comparison of TIDE spectra between leaves of the same T1 plant. Three true leaves (A, B and C) of a T1 plant were sampled. Genomic DNA was extracted, PCR amplified and sequenced. The indel spectrum is visualized with an estimated overall efficiency and the frequency of each indel using TIDE. Bars indicate the number of sequences with a given indel size. Pale green bars (indel size of zero) represents WT or base substitution alleles. A, example of a highly efficient edited T1 plant with low chimerism. B, example of a highly efficient edited T1 plant with high chimerism. C, example of a T1 plant with low editing efficiency.

All but one sgRNA had high editing efficiencies with the median efficiency being higher than 80% (Figure 2A). Notably, VQ33-38, the sgRNA predicted by all three algorithms to have the worst efficiency (Table S2) had one of the highest efficiencies *in planta*. Next, we used the data generated, to investigate chimerism in the T1 plants. The most frequently observed mutation is a 1 bp insertion, followed by deletions of increasing size (Figure 2B). Large insertions were very uncommon. However, depending on the sgRNA larger deletions of a particular size were often observed. Potentially this is related to MMEJ, whereby regions of ≥ 3 bp microhomology help initiate polymerase Θ repair by annealing of single-stranded DNA overhangs (Black *et al.*, 2013, Shen *et al.*, 2017). In summary, we show high rates of CRISPR/Cas9 mutagenesis in Arabidopsis T1 somatic tissue for most tested sgRNAs and that TIDE is a robust method to evaluate sgRNA efficiency.

**Figure 2.**
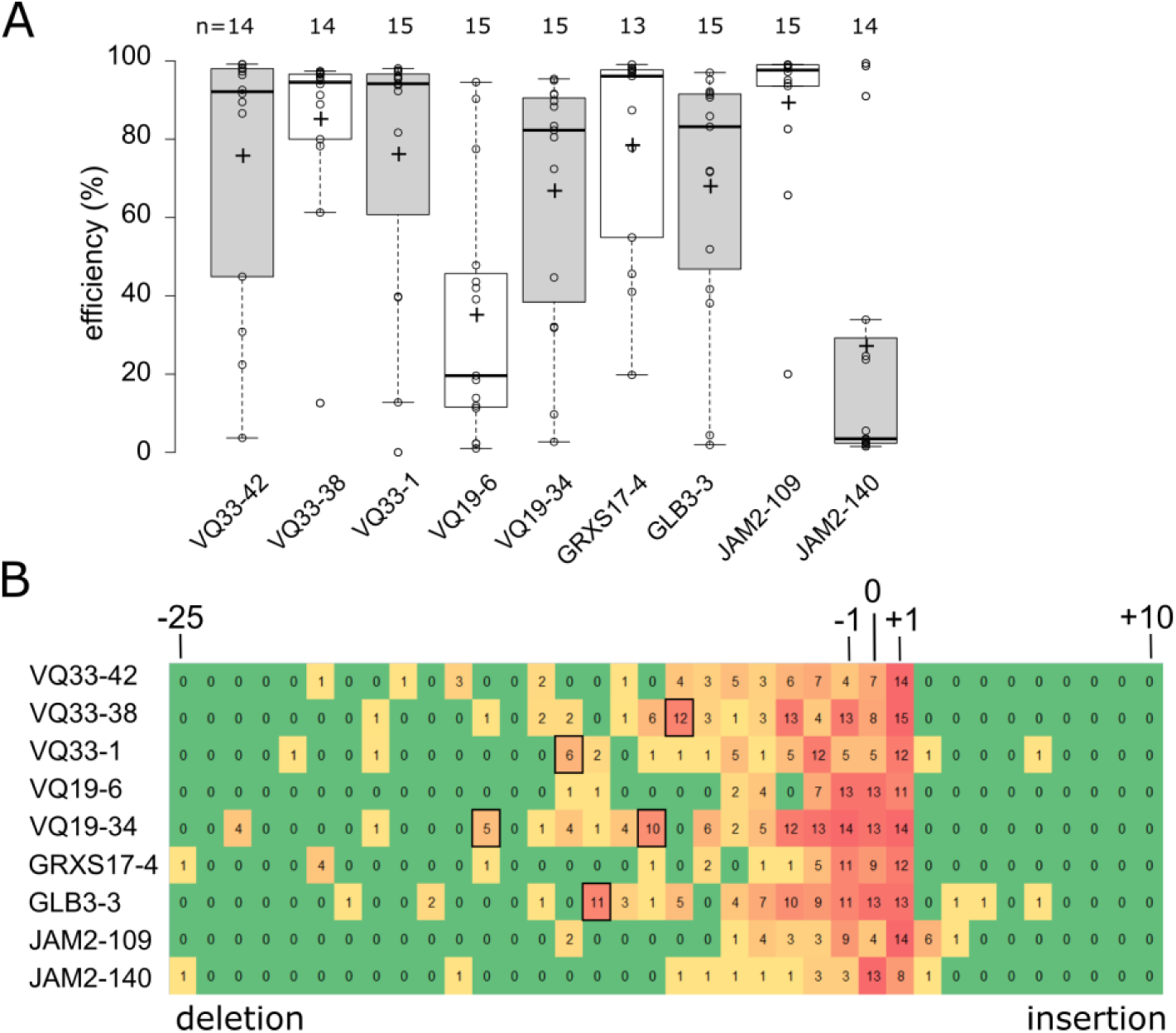
High gene editing efficiency in Arabidopsis T1 generation. A, boxplots showing TIDE estimated editing efficiencies for up to 15 T1 plants for nine different sgRNAs. +, mean; horizontal line, median; open circles, individual data points. B, heat map showing the number of T1 plants with at least 1% estimated frequency of an indel of a given size. Boxed are larger deletions observed in multiple T1 plants.

### Inheritance of mutations to T2

Focusing on *GLB3*, we investigated the heritability of these after selfing and selected for T2 progeny that had lost the T-DNA (Cas9 null-segregants). First, we identified three T1 lines with a single T-DNA locus by segregation analysis of the kanamycin resistance marker in T2 seedlings. Of these three lines we germinated 14 to 17 seedlings on soil, prepared genomic DNA and genotyped using Cas9 specific primers to identify null-segregants (Figure 3A). The genomic DNA of these plants was re-used to amplify the target site and sequencing data was analyzed using TIDE to identify genotypes at the target locus. All 15 tested null-segregants were found to be non-chimeric: 8 were WT, 5 heterozygous and 2 were homozygous. Hence, inherited mutations were present in the T2 progeny of all three independent T1 lines. Although we only detected the desired homo-allelic Cas9 null-segregants in the progeny of one T1 line, heterozygous alleles will lead to the desired genotypes in the next generation. An outcome also overrepresented in T1 somatic mutations for *GLB3,* and frequently observed in the inherited mutations from independent events was a 10 bp deletion (Figure 3B). Lastly, we identified a heritable T to A substitution which led to a single nucleotide variation (SNV) and here results in a premature stop codon (Figure 3B and 3C). This occurs when a single bp deletion is followed by a single bp insertion, an event very rarely observed for CRISPR/Cas9 (Kim *et al.*, 2017). In conclusion, the pDE-Cas9 vectors allow for efficient and inheritable genome editing in Arabidopsis with the possibility of producing transgene free homo-allelic mutants in the T2 generation.

**Figure 3.**
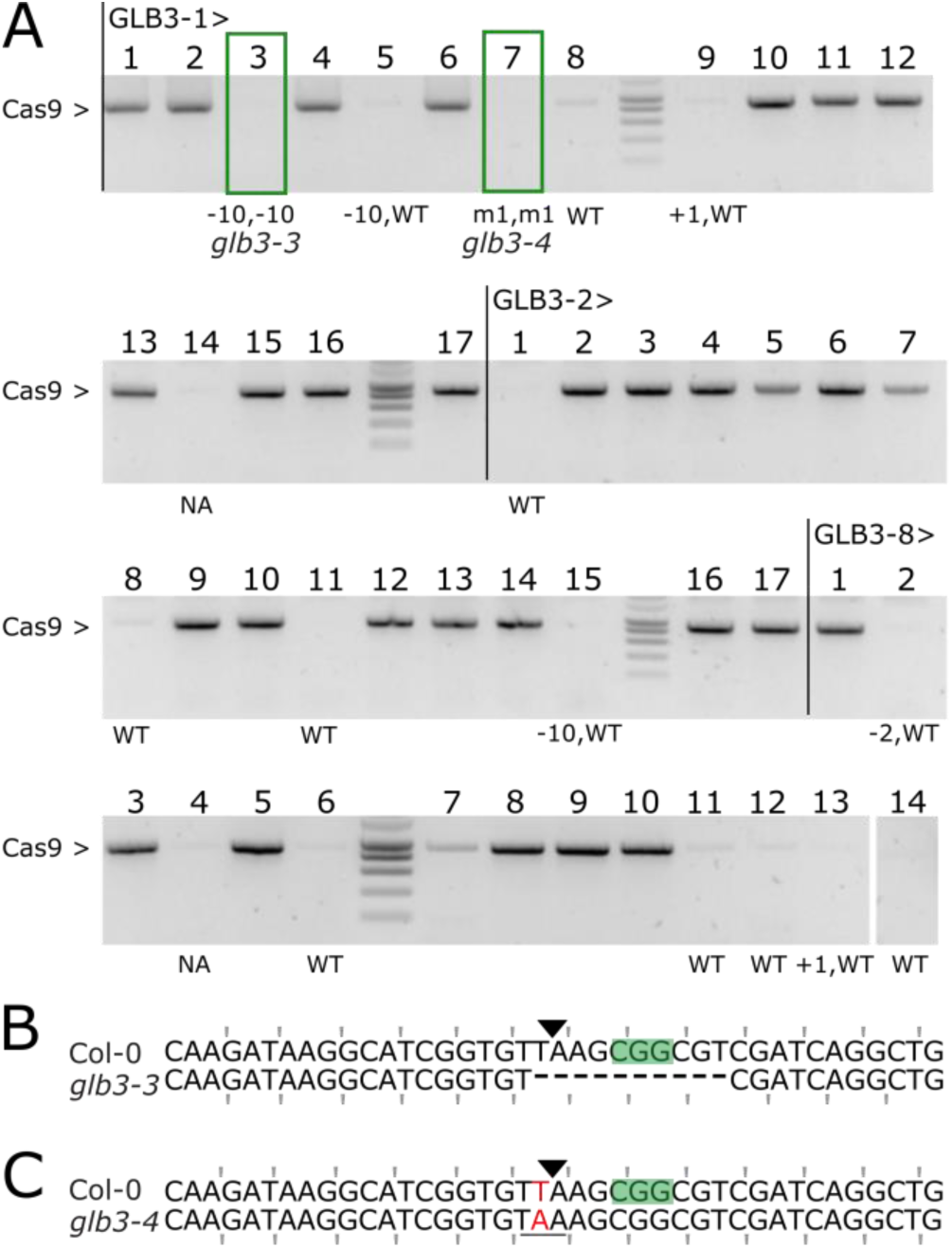
Inheritance of CRISPR/Cas9 mutations. A, PCR amplification of the Cas9 transgene in T2 seedlings from 3 independent GLB3 lines: -1, -2 and -8. Genotypes for all null-segregants were estimated using TIDE. NA, not assayed; WT, wild-type; m1, 1bp substitution. Boxed plants were continued. B-C, Sequence alignment of the targeted locus for Col-0 and glb3-3 (B, Line 1, plant 3) or glb3-4 (C, Line 1, plant 7). PAM is highlighted, the Cas9 cut site indicated with a triangle. Mutated bases are in red, deleted bases replaced by an en dash. The reading frame is marked.

**Table 1.**
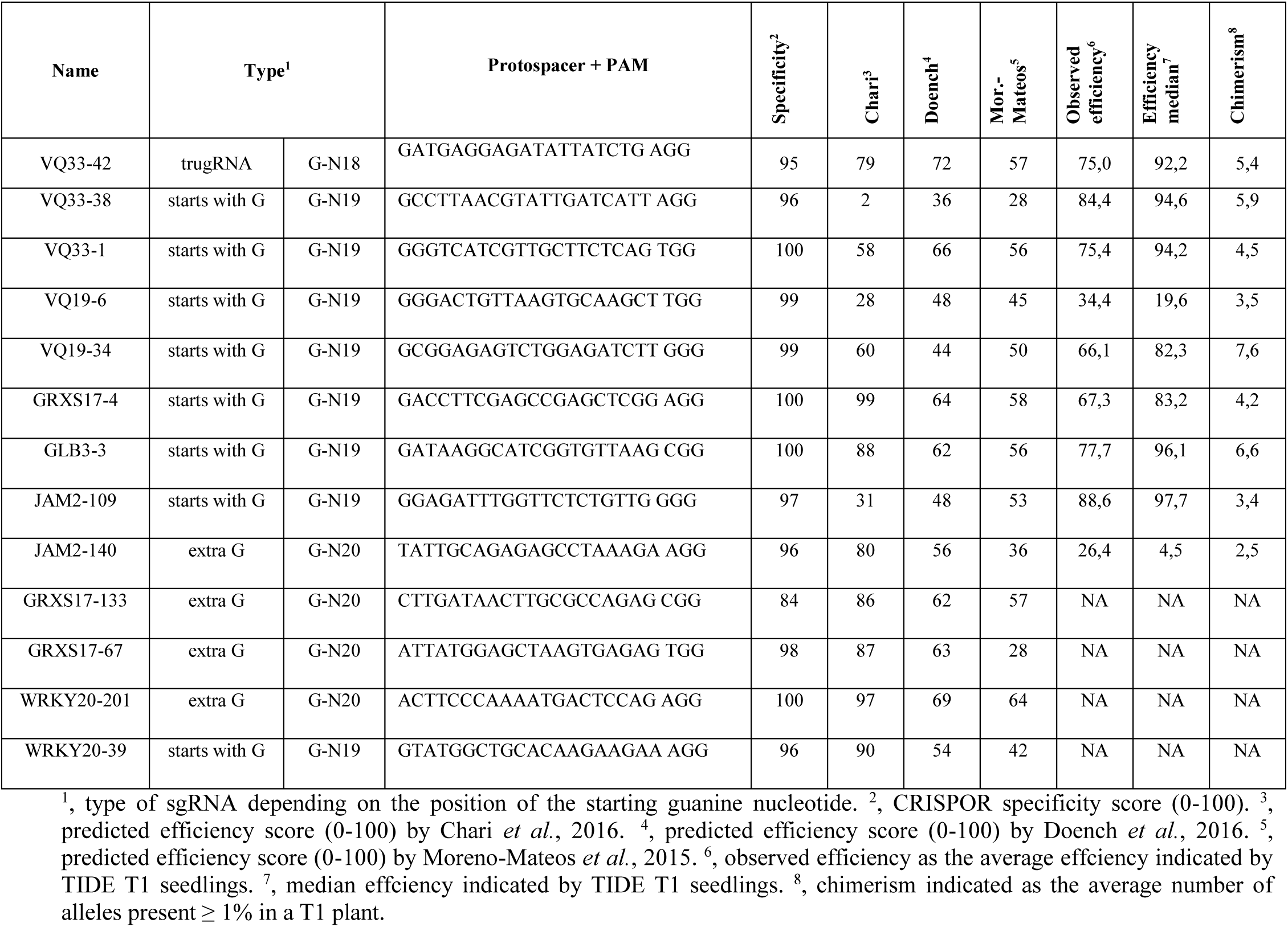
sgRNA parameters used in this study.

### Isolation of a new grxs17 CRISPR allele

Previously we characterized in detail two independent knock-out alleles of *GRXS17*, a gene encoding a component of the FeS cluster assembly pathway (Iñigo *et al.*, 2016). The allele *grxs17-1* (SALK_021301) contains a T-DNA in the second exon (Figure 4A), whereas the *grxs17-2* allele expresses an antisense construct (Cheng *et al.*, 2011).

**Figure 4.**
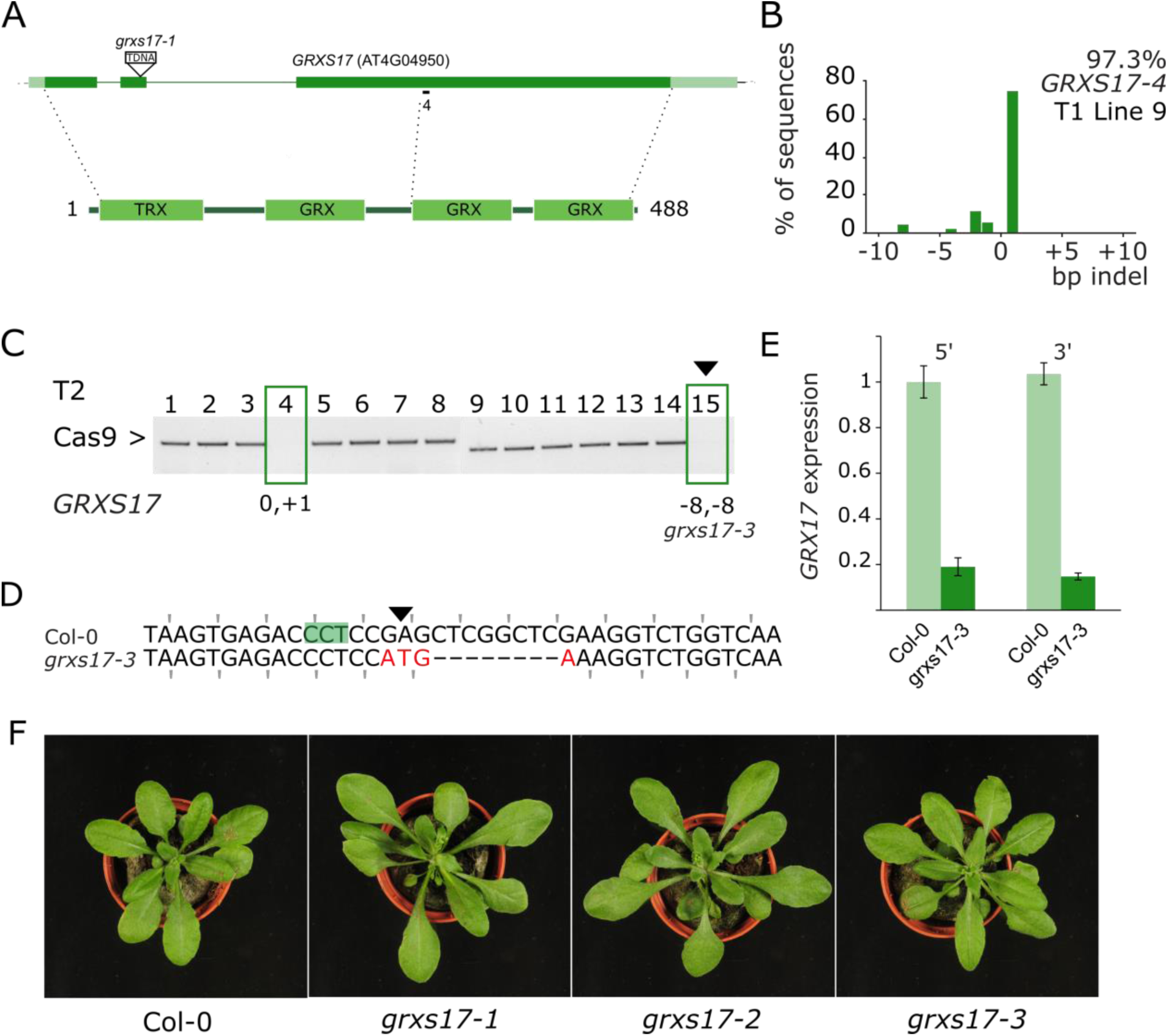
*grxs17-3* does not show a *grxs17-1* phenotype. A, Gene and protein structure of GRXS17. The location of the *grxs17-1* T-DNA insertion is indicated and the sgRNAs used in this study. Dark green boxes designate exons; light green boxes, UTRs; solid lines, introns. TRX, thioredoxin domain; GRX, glutaredoxin domain. B, TIDE analysis of T1 line 9. Genomic DNA was PCR amplified and sequenced. The indel spectrum is visualized with an estimated overall efficiency and the frequency of each indel. C, PCR amplification of the Cas9 transgene. Null-segregants are boxed and the continued plant marked with a triangle. TIDE estimated genotypes for *GRXS17* are given for the null segregants. D, Sequence alignment of the targeted locus for Col-0 and grxs17-3 (Line 9, plant 15). PAM is highlighted, the Cas9 cut site indicated with a triangle. Mutated bases are in red, deleted bases replaced by an en dash. The reading frame is marked. E, *GRXS17* gene expression analyzed by RT-qPCR. Expression relative to Col-0 is plotted using primers annealing both at the 5’and the 3’ of the transcript and the mutation. F, rosette phenotypes of Col-0, the T-DNA insertion line *grxs17-1*, the antisense line *grxs17-2* and the *grxs17-3* CRISPR allele.

A T1 parental line described above that showed high editing efficiency (97.3%) in somatic tissue and had a single T-DNA locus was identified (Figure 4B). Two Cas9 null-segregants of the T2 progeny were genotyped using TIDE (Figure 4C). This yielded the *grxs17-3* allele that was predicted to have a (-8,-8) genotype. Inspection of the sequence in T3 plants revealed an additional 4 bases mutated, nevertheless leading to loss of the reading frame (Figure 4D). Using RT-qPCR, we could observe strong downregulation (~80%) of the entire *GRXS17* transcript (Figure 4E). This is probably the result of nonsense-mediated decay (NMD), a process triggering mRNA degradation in case a premature stop codon is present (Popp and Maquat, 2016). However, as there is no exon-exon boundary 3’ of the premature stop codon, this can be a case of exon-junction complex (EJC)-independent NMD, wherein NMD is triggered by a long 3’ UTR (Fatscher *et al.*, 2014). Remarkably, the elongated leaf developmental phenotype present in both *grxs17-1* and *grxs17-2* was not visible in *grxs17-3* (Figure 4F). GRXS7 is a multidomain protein with an N-terminal thioredoxin (TRX) domain followed by three glutaredoxin (GRX) domains (Figure 4A). The human GRX3 ortholog has only 2 GRX domains, whereas the yeast Grx3/Grx4 orthologs have only one GRX domain (Couturier *et al.*, 2014). We hypothesize that the *grxs17-3* allele is not a null allele and possibly expresses a C-terminally truncated GRXS17 protein with a functional TRX and GRX domain.

### A dual sgRNA approach for gene deletions

Choice of the sgRNA target site is pivotal to generate a reliable knock-out. Genes can contain alternative start codons, have alternative first exon usage, exon skipping and/or C-terminally truncated proteins and therefore might still be partially functional as exemplified above. In-depth knowledge on the gene structure, transcript and protein is therefore advisable. However, in many cases this information is not complete. Therefore, we examined in Arabidopsis a dual sgRNA approach in which two sgRNAs target the same gene to remove a large part (Chen *et al.*, 2014, Zhang *et al.,* 2016, Ordon *et al.*, 2017).

Using a MultiSite Gateway based sgRNA multiplexing approach we previously described (Ritter *et al.*, 2017) we co-expressed two sgRNAs in pDE-Cas9Km. We used this method to target the gene encoding the transcription factor WRKY20, which is closely related to WRKY2, with two sgRNAs. For the latter, a characterized T-DNA insertion mutant *wrky2-1* is available representing a strong loss-of-function or null allele (Ueda *et al.*, 2011). We transformed the *wrky2-1* background with a dual sgRNA construct for *WRKY20*, predicted to remove a 247 bp fragment encoding the first WRKY protein domain in addition to putting the remainder of the sequence out of frame (Figure 5A). Without any phenotypic selection, we applied the same workflow as before. We selected four independent T1 lines showing high levels of the expected deletion and containing a single T-DNA locus (Figure 5B). For each line, one or more null-segregants were identified in T2 (Figure 5C) and genotyped for the *WRKY20* locus. Of seven Cas9 null-segregants successfully genotyped, two plants were homozygous for the expected deletion, three heterozygous and two wild-type (Figure 5D). Sequence analysis of two homozygous deletion mutants showed that *wrky2-1 wrky20-1* (plant A15-8) had the predicted 247 bp deletion, whereas the other allele *wrky2-1 wrky20-2* (plant B2-5) had only 246 bp deleted, possibly restoring the reading frame (Figure 5E). This shows that a dual sgRNA approach for deleting gene fragments is feasible with relatively few numbers of genotyped plants.

**Figure 5.**
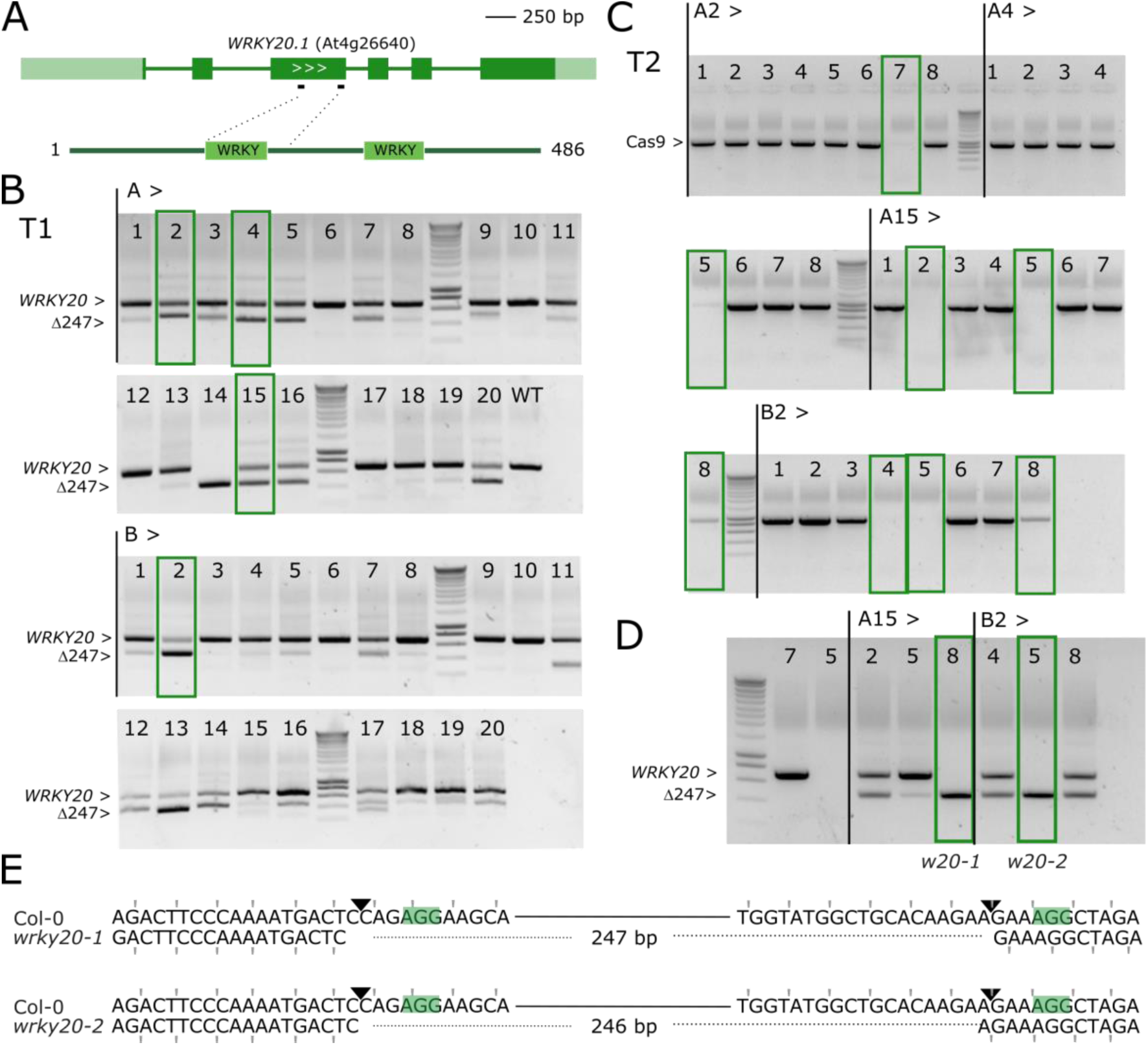
*WRKY20* dual sgRNA approach. A, genomic structure of *WRKY20* and location of the sgRNAs. Dark green boxes designate exons; light green boxes, UTRs; solid lines, introns. B, PCR analysis of T1 lines. Leaf genomic DNA of 2 batches (A and B) of 20 chimeric T1 plants was PCR amplified. The expected size of the WT *WRKY20* amplicon is indicated as well as the expected size of the deletion of 247 bp between Cas9 cut sites. Four continued T1 lines having one T-DNA locus are highlighted with green boxes. C, Cas9 PCR for the four continued lines in T2 generation. Putative Cas9 null-segregants are indicated with green boxes. D, Cas9 null-segregants were genotypes for WRKY20. The selected lines A15-8 (*wrky2-1 wrky20-1*) and B2-5 (*wrky2-1 wrky20-2*) are boxed. E, Sequence alignment of the simultaneously targeted loci for Col-0 and alleles *wrky20-1* and *wrky20-2*. PAMs are highlighted, the Cas9 cut sites indicated with triangles. Deleted bases are indicated with dashed lines. The reading frame is marked.

Next, we combined two sgRNAs targeting *VQ33* (VQ33-42 and VQ33-1) that displayed high efficiency when tested individually (Figure 2). Working together, they are predicted to remove a fragment of 459 bp, virtually removing the *VQ33* coding sequence (Figure S3A). We proceeded with the same workflow as for *WRKY20* (Figure S3B-D). Out of four Cas9 null-segregants, two were homozygous for the expected gene fragment deletion, one heterozyogous and one WT. The allele *vq33-1* (plant 11-8), albeit it had an extra 1 bp insertion, still led to a 458 bp out-of-frame deletion (Figure S3E).

In summary, we established a straightforward dual sgRNA approach to obtain plants homozygous for relatively large deletions of gene fragments in the T2 generation in *Arabidopsis thaliana*.

**Figure S3.**
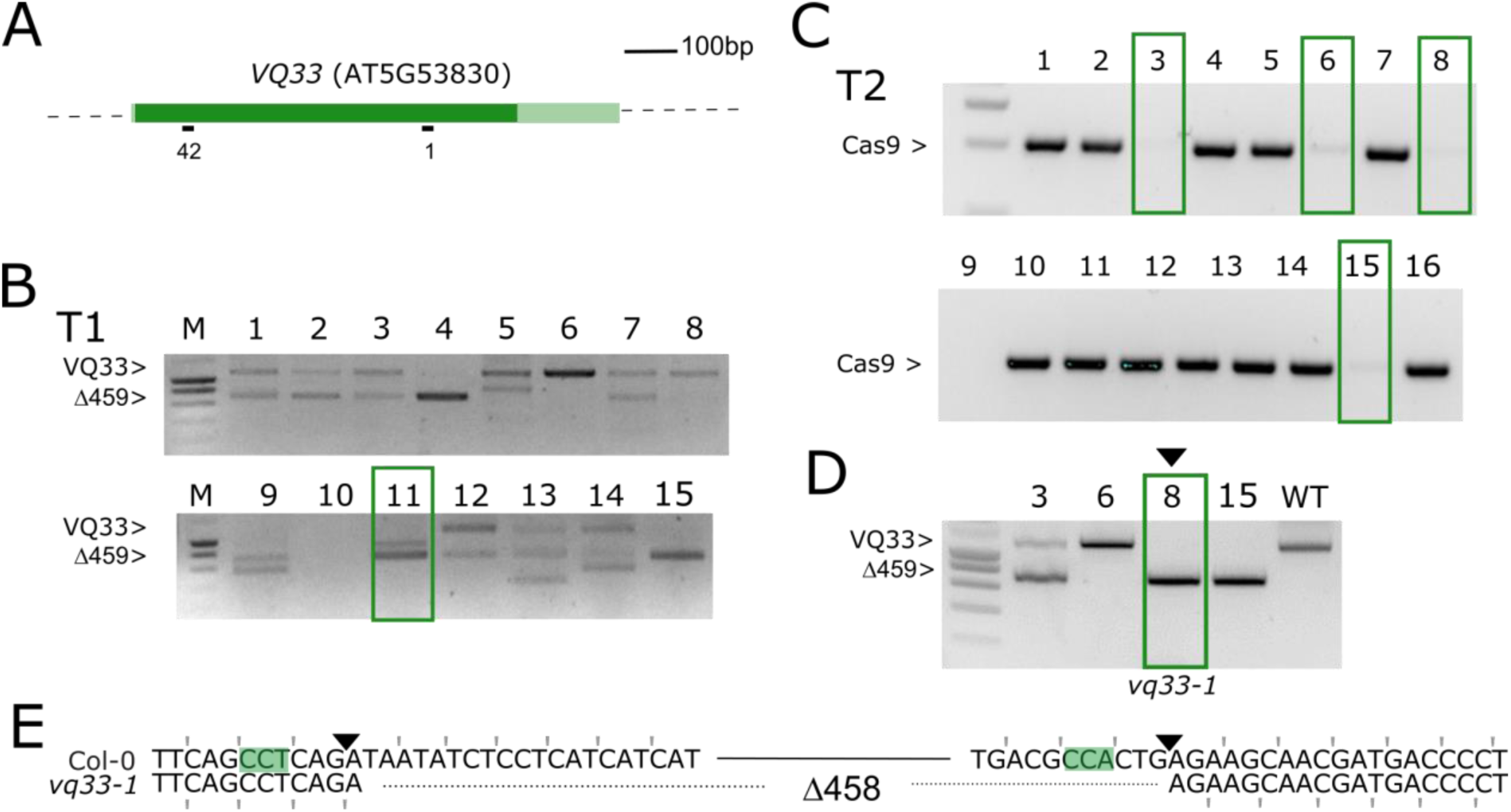
*VQ33* dual sgRNA approach. A, genomic structure of *VQ33* and location of the sgRNAs. Dark green boxes designate exons; light green boxes, UTRs; solid lines, introns. B, PCR analysis of T1 lines. Leaf genomic DNA of 16 chimeric T1 plants was PCR amplified. The expected size of the WT *VQ33* amplicon is indicated as well as the expected size of the deletion of 459 bp between Cas9 cut sites. One T1 line having one T-DNA locus that was continued is highlighted with a green box. C, Cas9 PCR for the continued line in T2 generation. Putative Cas9 null-segregants are indicated with green boxes. D, Cas9 null-segregants were genotypes for *VQ33*. The selected plant 11-8 (*vq33-1*) is indicated with a triangle. E, Sequence alignment of the simultaneously targeted loci for Col-0 and *vq33-1*. PAMs are highlighted, the Cas9 cut sites indicated with triangles. A 458 bp deletion was detected and is indicated with dashed lines. The reading frame is marked.

### grxs17-4 confirms the grxs17-1 developmental phenotype

Next, we tried the dual sgRNA approach for *GRXS17*. We targeted the first sgRNA (GRXS17-133) at the 5’ end and the second sgRNA (GRXS17-67) at the 3’ end of the gene to remove 1953 bp and *GRXS17* almost entirely (Figure 6A). The *GRXS17* locus was amplified for sixteen independent T1 plants using primers spanning the expected deletion. In comparison with *VQ33* and *WRKY20*, only two plants clearly showed bands of the expected size for the predicted deletion (Figure 6B). Two identified Cas9-null segregants (Figure 6C) did not show the expected large deletion, but instead an indel was found at the first sgRNA site in the first exon leading to a frameshift (Figure 6D). We re-named this allele *grxs17-4*. Confirming our hypothesis that *grxs17-3* is indeed not a null allele, *grxs17-4* showed the leaf phenotype of *grxs17-1* and *grxs17-2* (Iñigo *et al.*, 2016, Figure 6E). In conclusion, in the event the dual sgRNA approach does not yield the designed gene fragment deletion, each individual sgRNA may lead to useful alleles.

**Figure 6.**
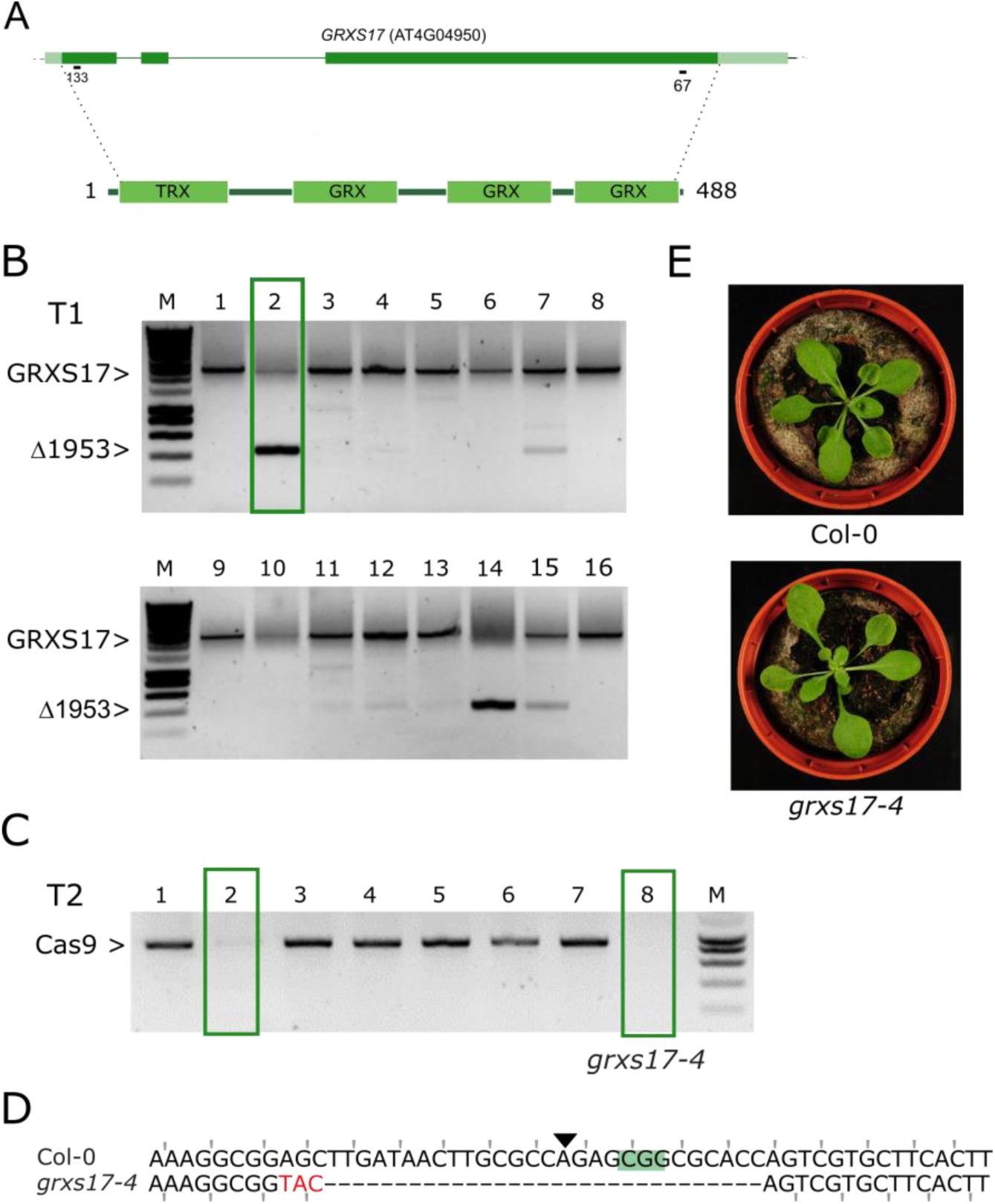
A dual sgRNA approach for *GRXS17.* A, genomic structure of *GRXS17* and location of the sgRNAs. Dark green boxes designate exons; light green boxes, UTRs; solid lines, introns. B, PCR analysis of T1 lines. Leaf genomic DNA of 16 chimeric T1 plants was PCR amplified. The expected size of the WT *GRXS17* amplicon is indicated as well as the expected size of the deletion of 1953 bp between Cas9 cut sites. One T1 line having one T-DNA locus that was continued is highlighted with a green box. C, Cas9 PCR for 8 T2 CRISPR plants. Putative Cas9 null-segregants are indicated with green boxes. D, Sequence alignment of the sequence surrounding the 5’ sgRNA site for Col-0 and *grxs17-4* (Line 2, plant 8). PAM is highlighted, the Cas9 cut site indicated with a triangle. Mutated bases are in red, deleted bases replaced by an en dash. The reading frame is marked. E, representative rosette phenotypes of WT Col-0 (top) and *grxs17-4* (bottom).

## DISCUSSION

### Efficient CRISPR/Cas9 gene editing in Arabidopsis

The CRISPR/Cas9 technology shows promise to speed up reverse genetics experiments in Arabidopsis. Here we demonstrate efficient recovery of Cas9-free Arabidopsis mutants using single and double sgRNA constructs in the T2 generation without phenotypic selection. Previous negative experiences with CRISPR/Cas9 have been attributed to the weak activity of the 35S promoter in germ-line cells (Wang *et al.*, 2015). The promoter used here, PcUBI, is expressed widely, but detailed expression in germ-line cells has not yet been studied (Kawalleck *et al.*, 1993). Other vector elements have been reported to play a role such as the vector backbone (Mao *et al.*, 2016), Cas9 coding sequence (Johnson *et al.*, 2015) and the terminator sequence (Wang *et al.*, 2015). We did not observe any obvious differences using either *nptII* or *bar* as selection markers. Systematic analysis of all vector parameters is now achievable using modular cloning systems, which might allow identification of the best combinations (Vazquez-Vilar *et al.*, 2016).

We consider the workflow presented here as already an acceptable workload comparable to the routine generation of overexpression lines. Nonetheless, several improvements have recently been developed. For example, a fluorescent marker for identification of transgenic T1 seeds has been reported (Tsutsui and Higashiyama, 2016) and also recently cloned into pDE-Cas9 for CRISPR/Cas9 in *Camelina sativa* (Morineau *et al.*, 2016). When Cas9 is driven with a promoter active in the egg cell, non-chimeric homozygous or bi-allelic mutants can already be retrieved in the T1 generation, although Cas9 null-segregants also only appear in T2 (Wang *et al.*, 2015, Yan *et al.*, 2016, Mao *et al.*, 2016, Eid et al;, 2017).

### TIDE as a useful tool to study mutations

Efficiency of CRISPR/Cas9 also clearly depends on the choice of sgRNA, although all sgRNAs tested in this study were active to some degree. Several models have been constructed to predict on-target editing efficiency based on the sgRNA primary sequence and on-target efficiency data from metazoans (Doench *et al.*, 2016, Moreno-Mateos *et al.*, 2015). Due to the lack of sufficient data, no plant-specific design models are currently available. As previously reported (Ordon *et al.*, 2016), we did not observe any obvious correlation between these predictions and our observed efficiencies in Arabidopsis. It is unclear why this is the case for a heterologous system such as CRISPR/Cas9. Therefore - for the time being - we continue to take into account metazoan models when designing plant sgRNAs. It has been suggested to pre-screen sgRNAs in protoplasts (Li *et al.*, 2014). Given the ease of Arabidopsis transformation via floral dip, we conclude from this study that designing several sgRNAs for the same target and testing somatic mutations in T1 might be an equally rapid method to identify efficient sgRNAs, while simultaneously obtaining the desired mutants.

Several methods have been used to study CRISPR/Cas9-induced mutations, most importantly cleaved amplified polymorphic sequence (CAPS), T7 endonuclease, next-generation sequencing and high-resolution melting curve analysis (Denbow *et al.*, 2017). The method used here, TIDE (Brinkman *et al.,* 2015), has several advantages. First, it does not require a restriction enzyme site overlapping the Cas9 cut site as in CAPS. Secondly, it allows the starting genomic DNA to be relatively impure allowing for more economic DNA extraction methods compared to T7-based assays. Thirdly, it uses standard capillary Sanger sequencing that can be readily performed for even a single sample. Fourthly, it can provide an insight in the indel spectrum of mosaics similar to next-generation sequencing as well as providing an idea of overall efficiencies. These TIDE efficiencies are likely an underestimation. For example, TIDE is unable to detect rare SNVs as observed for *glb3-4*. The *grxs17-3* allele also revealed that mutations can be more complex than predicted by TIDE: a predicted 8 bp deletion was actually a 12 bp deletion combined with a 4 bp insertion.

### Know your target gene

The absence of the typical *grxs17* phenotype in the CRISPR allele *grxs17-3* is an example of how it is important to study independent alleles made with either different sgRNAs or with other methods when interpreting phenotypes of CRISPR/Cas9-generated alleles as knock-out effects. When sufficient information is available, especially on alternative transcripts and protein domain structures, sgRNA target sites can be chosen to maximize the chance of a complete knock-out as a result of an indel mutation at that site. Additionally, one may disrupt the gene more dramatically by removing a larger gene fragment using a dual sgRNA approach. The use of CRISPR/Cas9 for gene deletion has been pioneered in mammalian systems (Chen *et al.*, 2014, Zhou *et al.*, 2014, Ran *et al.*, 2013, Canver *et al.*, 2014). In Arabidopsis, a dual sgRNA approach for gene deletion was reported by Zhao *et al.*, 2016 and Ordon *et al.* 2017. In Zhao *et al.*, homozygous deletion mutants were obtained for the *AtMIR827a* and *AtMIR169a* loci in the T2 or T3 generation, respectively. The size of the deletion and efficiency seem to correlate inversely in mammalian cells (Canver *et al.*, 2014) and plants (Ordon *et al.*, 2016). Similarly, when attempting to cut out a 1953 kb fragment in *GRXS17*, it failed to be inherited, while clearly being present in T1 somatic cells. In contrast, 247 bp and 459 bp fragment deletions were easily obtainable for *WRKY20* and *VQ33* respectively. Therefore, while deleting whole genes might be tempting, it is more practical targeting genes with two sgRNAs in the 5’ coding sequence. This has the additional advantage, that when one sgRNA has a low efficiency, the construct will still yield potential knock-out mutations at the other sgRNA site. It has been proposed from work in tomato protoplasts that in most cases when a single sgRNA is used, NHEJ results in perfect repair and therefore using two sgRNAs could be more efficient to obtain mutants (Čermák *et al.*, 2017). Finally, the double-sgRNA approach has an advantage of easy visual genotyping of mutants based on amplicon lengths.

### New alleles for GRXS17

*GRXS17* encodes the Arabidopsis ortholog of human *GRX3/PICOT* and yeast *Grx3/Grx4*. Although a role for GRXS17 in iron-sulfur cluster assembly is conserved in all of these organisms, plant-specific functions for GRXS17 are apparent (Iñigo *et al.*, 2016, Kneustig *et al.*, 2016). Interestingly, AtGRXS17, HsGRX3 and ScGrx3/4 differ in the number of GRX domains that are C-terminal of the TRX domain with three, two and one domain present, respectively. The new *grxs17-3* allele presented here might have residual expression of a truncated GRXS17 with only one GRX domain — similar to ScGrx3/4 — and could therefore be helpful in studying plant-specific GRXS17 roles.

## MATERIALS AND METHODS

### Design of sgRNAs

In general, sgRNAs were selected for specificity using CRISPR-P (http://cbi.hzau.edu.cn/cgi-bin/CRISPR, Lei *et al.*, 2014), taking into account predicted on-target efficiencies using sgRNAscorer (https://crispr.med.harvard.edu/sgRNAScorer/, Chari *et al.*, 2015). An updated overview of estimated sgRNA parameters by CRISP-OR (http://crispor.tefor.net/, Haeussler *et al.*, 2016) can be found in Table 1.

### Cloning of CRISPR/Cas9 constructs

CRISPR/Cas9 constructs were cloned as previously described (Figure S1, Fauser *et al.*, 2014, Ritter *et al.*, 2017). Briefly, for each guide sequence, two complementary oligos with 4bp overhangs (Supplementary Table S1) were annealed and inserted via a cut-ligation reaction with BbsI (Thermo) and T4 DNA ligase (Thermo) in a Gateway ENTRY sgRNA shuttle vector. This is either pEN-C1.1 (Fauser *et al.*, 2014) for single sgRNA constructs, or pMR217 (L1-R5) and pMR218 (L5-L2) (Ritter *et al.*, 2017) for the dual sgRNA approach. The 5’ overhang already contains the G initiation nucleotide of the AtU6-26 polIII promoter. Next, using a Gateway LR reaction (ThermoFisher), one or two sgRNA modules were then combined with pDE-Cas9 (Basta, Fauser *et* al, 2014) or pDE-Cas9Km (pMR278, Ritter *et al.*, 2017) to yield the final expression clone.

### Plant transformation

Expression clones were introduced in the Agrobacterium strain C58C1 (pMP90) using electroporation, which was used to transform Arabidopsis using the floral dip method (Clough and Bent, 1998).

### Plant Material and Growth Conditions

*Arabidopsis thaliana* Col-0 were grown at 21°C under long day (16-h light/8-h dark) conditions. Rapid selection of seeds with kanamycin and phosphinothricin (BASTA^™^) selection was performed as described (Harrison *et al.*, 2006).

### Selection of CRISPR/Cas9 mutants

A scheme of our strategy is given as Figure S4. Typically, 16 kanamycin-or BASTA-resistant T1 plants are selected *in vitro* and transferred to a growth room. After 14 days, a single leaf is harvested and genomic DNA prepared using Edwards buffer (Edwards *et al.*, 1991). Next, 5 μl template gDNA was used as a template in a standard 20 μl volume PCR reaction using GoTaq^®^ (Promega) with the supplied Green GoTaq® Reaction Buffer. For single sgRNA constructs, the amplicon was treated with ExoSAP-IT^™^ (Thermo) and sequenced by standard capillary sequencing at the VIB Genomics Core Facility (https://corefacilities.vib.be/gsf). Quantitative sequence trace data was decomposed using TIDE (https://tide.nki.nl/) using standard settings, except for the indel size range, which was set on the maximum (50). Primers for TIDE were designed using Primer3 (http://bioinfo.ut.ee/primer3-0.4.0/) using standard parameters. Approximately 700 bp asymmetrically surrounding the Cas9 cut site was amplified. The amplification primer at 200 bp from the site was used for sequencing.

For each independent T1 line, approximately 64 T2 seeds were selected on either BASTA or kanamycin. Resistant versus sensitive seedlings were analyzed using a chi-squared test and lines presumably having a single T-DNA locus continued. Typically, 15 seedlings of the most promising line (highest T1 efficiency, expected segregation) were grown on non-selective media and genotyped for the presence of the T-DNA locus using Cas9-specific primers (Table S1). Cas9 null-segregants are then analyzed for modifications at the locus of interest. The most promising plants are then propagate to T3, in which absence of Cas9 and presence of the mutation/deletion is confirmed by PCR and sequencing.

**Figure S4.**
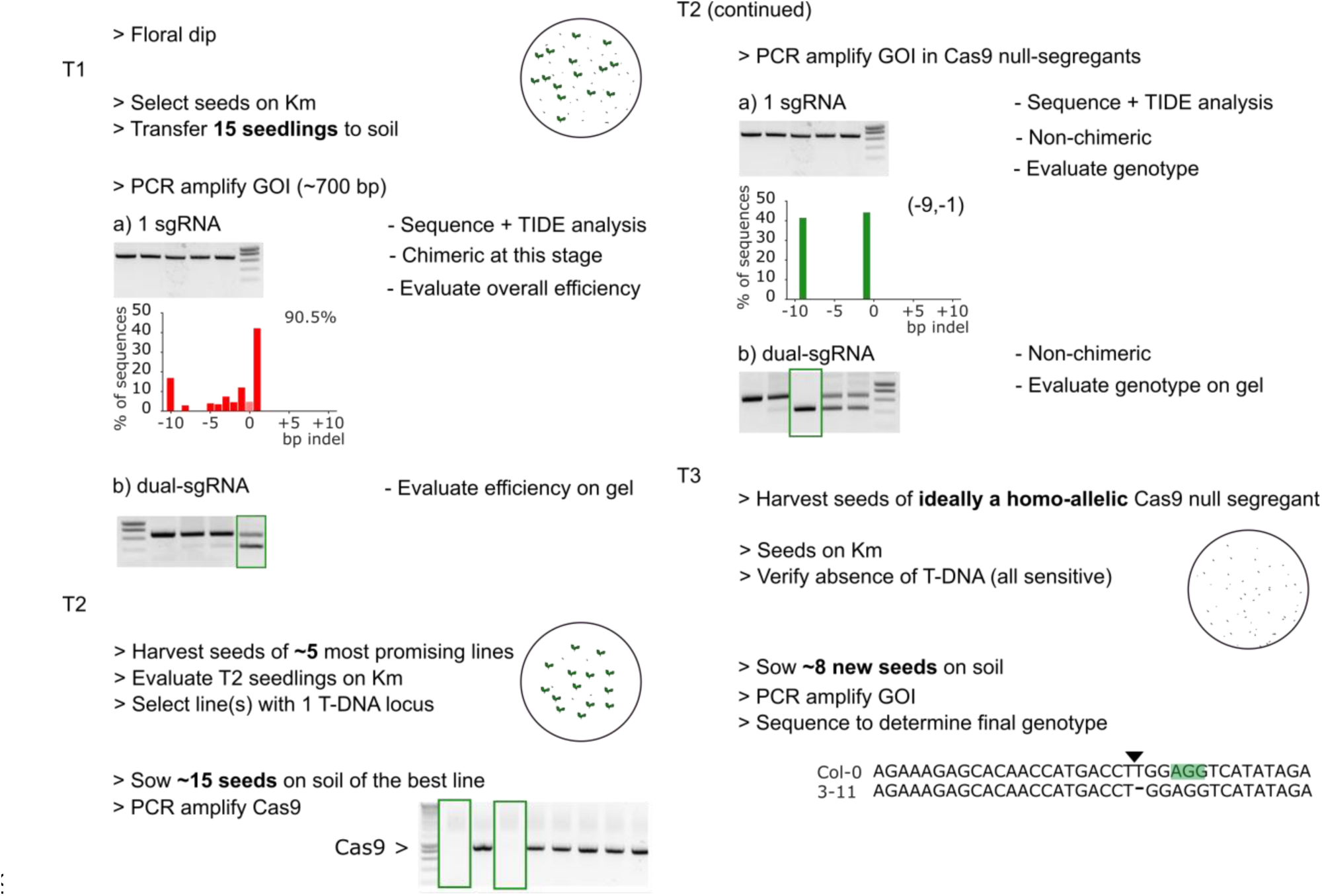
*CRISPR workflow*. Schematic overview of the selection of Cas9 null-segregants with bi-allelic mutations in the T2 generation.

### Amplicon subcloning

For confirmation of TIDE spectra, the PCR amplicon was cut from gel, purified using GeneJET PCR purification kit (Thermo Scientific) and cloned into pJET1.2 using the CloneJET PCR cloning kit (Thermo Scientific). Individual clones were sequenced using capillary electrophoresis.

### RT-qPCR

Seedlings were grown in the same conditions as in Iñigo *et al.*, 2016. Seedlings were frozen in liquid nitrogen and total RNA was extracted using RNeasy plant mini kit (Qiagen) and DNAse I (Promega) treatment. Next, 1 μg of RNA was used for cDNA synthesis using iScript kit (Bio-Rad). qRT-PCR was performed on a LightCycler 480 system (Roche) using the Fast Start SYBR Green I PCR mix (Roche) with three biological repeats and three technical repeats. Data were analyzed using the second derivative maximum method and relative expression levels were determined using the comparative cycle threshold method. Primer sequences are provided in Supplemental Table S1.

## ACCESSION NUMBERS

Accession numbers of the genes used in this study: *GRXS17*, AT4G04950; *VQ19/MVQ4*, AT3G15300; *VQ33/MVQ3*, AT5G53830; *WRKY20*, AT4G26640; *WRKY2*, AT5G56270; *JAM2/bHLH13*, AT1G01260; *GLB3*, AT4G32690. T-DNA lines used: *grxs17-1*, SALK_021301; *wrky2-1*, SALK_020399.

## SUPPLEMENTAL DATA

Figure S1. Cloning procedures and vector maps.

Figure S2. Comparison of TIDE spectra between leaves of the same T1 plant.

Figure S3. *VQ33* dual sgRNA approach.

Figure S4. CRISPR workflow.

Table S1. Oligonucleotides used in this study.

## ACKNOWLEDGMENTS

We thank Carina Braeckman for Arabidopsis floral dip transformations, Karel Spruyt for photography, Wilson Ardiles Diaz for help with sequencing, Annick Bleys for help in preparing the manuscript, Holger Puchta for providing the pDE-Cas9 vector and Thomas Laux for providing *wrky2-1* seeds. We thank Lieven De Veylder, Thomas Jacobs and Ward Decaestecker for their helpful comments on the manuscript.

## AUTHOR CONTRIBUTIONS

L.P., A.B. and A.G. designed the research; L.P., R.D.C, S.I., J.G., C.W. and M.R. performed research; L.P. wrote the paper with help from all authors.

